# Unravelling the universal spatial properties of coral reefs

**DOI:** 10.1101/2024.11.14.623536

**Authors:** Àlex Giménez-Romero, Manuel A. Matías, Carlos M. Duarte

## Abstract

Coral reefs are under rapid decline due to human pressures such as climate change. Achieving the Kunming-Montreal Global Biodiversity Framework goals, which include restoring 30% of degraded habitats like coral reefs by 2030, requires a comprehensive understanding of their extent and structure, which has been hitherto lacking. We address this limitation based on the unprecedented canonical inventory of coral reefs extracted from the Allen Coral Atlas of shallow-water tropical reefs. We identified a total of 1,579,772 individual reefs globally, extending over a total of 52,423 *km*^2^ of ocean area with mean and median sizes of 3.32 ha and 0.3 ha, respectively. We unravelled three universal laws that are common to all coral reef provinces: the size-frequency distribution, the inter-reef distance distribution and the area-perimeter relation, which follow power laws with an exponent of 1.8, 2.33 and 1.26, respectively. We demonstrate that coral reefs develop universal fractal patterns characterised by a perimeter fractal dimension of *D*_*P*_ = 1.3 and a surface fractal dimension of *D*_*A*_ = 1.6. Our analysis shows that coral reefs display intricate fractal-like geometries and exhibit universal macroecological patterns, largely independent of their geographical location. The universality of the observed patterns suggests that these features possibly stem from the highly conserved interactions of biological, physical, and chemical processes. Over geological scales, these processes lead to reef landscape patterns common among all provinces, providing new information relevant to reef growth modelling.

## 1 Introduction

Coral reefs form some of the largest biogenic structures in the biosphere [1] and are prevalent in tropical coastal waters. Reefs often form complex and labyrinthine structures that protect the shorelines of tropical coastal nations while supporting biodiversity and providing food supply to local communities [2]. The formation of coral reefs has intrigued scientists for a long time, with Darwin formulating a model for oceanic atolls based on coral reefs accreting on volcanic structures [3]. Darwin’s model continues to stir the discussion [4] because it does not explain the diverse types of reef formations that are present in nature [5]. Coral reefs can form isolated, oval, or ring-shaped structures such as coral atolls; linear structures parallel to the shoreline, such as fringing or barrier reefs; and convoluted or highly branched structures (e.g., [6]). They also often present nested structures, such as smaller reefs within large coral reef lagoons [7]. A comprehensive model that accounts for this diversity of forms and configurations has yet to be developed.

Previous studies have examined the geometry of coral reefs [6–11], providing evidence of fractality at both individual reef [7, 11] and regional scales [6, 8]. However, a worldwide assessment of coral reef size and geometry has been limited by the lack of comprehensive data on reef form and size at a global scale. Accurately determining the size and geometry of coral reefs has become a pressing issue, as corals are rapidly declining, with an estimated 50% of coral cover lost worldwide from 1957-2007 [12]. According to the IPCC, if global warming reaches between 1.5 and 2.0 ºC above preindustrial levels, we might see a reduction in coral cover by 70% to 99% respectively [13]. In contrast, the Kunming-Montreal Biodiversity Framework [14] calls for stopping all biodiversity losses and restoring 30% of degraded habitats, including coral reefs, by 2030. This requires individual reef interventions and a thorough understanding of the form and size of coral reefs as an underpinning for the necessary conservation actions.

The release of the Allen Coral Atlas (ACA) [15], a worldwide mapping initiative that provides benthic habitat data of shallow-water (above 10 m deep) tropical reefs, presents an unprecedented opportunity to characterise the form and size of coral reefs on a global scale. This initiative surpasses prior efforts, such as NOAA’s Coral Reef Information System [16], Khaled bin Sultan Living Oceans Foundation [17], or Millennium Coral Reefs [18], by combining extensive global coverage of reef areas with the direct availability of processed habitat classifications obtained from recent high resolution (3 m) remote sensing data using deep learning models [19, 20]. This resource offers a unique vantage point for researchers, scientists, and conservationists to explore and understand coral reef ecosystems in novel and highly detailed ways that previous methods could not achieve. For instance, a recent study has already taken advantage of the ACA to assess the impact of global-scale biogeographical and evolutionary histories on coral reef habitats [21].

However, extracting the size and geometry of the individual coral reefs from the ACA is not straightforward. The ACA provides an interface that shows the classified benthic cover from satellite images and allows the user to retrieve the total area by cover class across an entire province or user-defined areas. Registered users can also download a georeferenced vector file that contains polygons of different cover classes. Some of these polygons represent single reefs, while others are partial components of larger individual reefs. Hence, while the ACA allows quantification of the area of each class at the province or user-defined levels, it was not designed to support the analysis of the shape and size of individual reefs. Measuring the spatial features of individual coral reefs requires significant post-processing of the data and enough computational capacity, which likely explains the limited use of the ACA in addressing current knowledge gaps concerning coral reef shape and size.

Here, we process data from the ACA’s 2022 release to compile a comprehensive global inventory of 1, 579, 772 individual shallow-water tropical coral reefs (>1000 m^2^) [22]. This dataset enables us to analyse several spatial properties of individual reefs, unravelling key macroecological patterns [23] of reef size and geometry, such as a universal power-law size-frequency distribution and constant fractal dimensions consistent across coral provinces. We also examined the relationships between the size, shape, and fractal geometry of coral reefs to identify distinctive patterns at the reef, province, and global levels. Our analyses improve and deepen our understanding of these complex ecosystems and provide elements to quantify the scale of the effort required to conserve and restore them. Open questions about the mechanisms behind the formation of the observed patterns suggest potential avenues for future research.

## 2 Methods

### 2.1 Global coral reef data

Global reef-mapping system data were obtained from the Allen Coral Atlas (ACA) [15], a publicly available dataset of high-resolution satellite imagery (2018-2020) and machine learning-based coral reef classifications [19, 20, 24]. We downloaded the data from the ACA website, which is already divided into the different coral reef provinces. The provinces mapped by the ACA reflect established patterns of coral reef biogeography with similar reef type and environmental conditions [15, 25]. The downloaded dataset consists of GeoJSON files for each coral province with several Polygons and Multiploygons forming the different benthic classes, including coral/algae, seagrass, microalgal mats, rock, rubble and sand [15].

The “coral/algae” class provides confidence that living corals may be present, so we selected this class to avoid false negatives for habitats classified as sandy, algal mats, rocky or rubble that are unlikely to have contained living corals within the limits of the resolution of the ACA at the time the images were retrieved. This was guided by the emphasis on coral cover in assessments of the health status, past, present, and future, of corals [26–28], best supported by the coral/algae category of the ACA. We acknowledge that this typology is rather limited compared to richer coral reef classification frameworks that have been proposed [20] but have not yet been adopted by the ACA.

We used the polygons classified as coral/algae to identify the calcifying community of coral reefs, and hereafter refer to this class as just coral reefs or reefs. This habitat class is characterised by a hard underlying framework with a benthic covering of coral and/or algae. The benthic cover of coral or algae should be at least 1%, usually more than 5% and sometimes exceed 40%, but it does not necessarily have a dominance of any of these groups over non-living substrate [15, 20]. With an average coral cover of 10-20% worldwide, most reef habitats, even those supporting extensive coral growth, are unlikely to be quantitatively dominated by living coral [20, 27]. Because our data is based on the coral/algae class of the ACA, we exclude reef structures of possible biogenic origin that did not support coral or algae at the time the images were retrieved, but that may have been covered by coral in the past. Despite the “rock” class could have been included to take this into account, this could also bias the results by including igneous rock outcrops that may not be or may not have originated from coral reef calcifying communities.

Even though the data are already provided in vector format, the reefs are not identified as separate entities, i.e., a single reef can be formed by many polygons or multi-polygon objects. Thus, we processed the dataset using the methods explained below to obtain a representation of individual reefs. We have used a global projected coordinate system covering 86°S to 86°N with meter units (Global, Equal-Area; EASE-Grid 2.0, EPSG:6933) for all our computations.

### 2.2 Coral reefs as clusters of connected coral/algae class polygons

A label assignment algorithm was developed to identify the different independent (not connected) components that make up coral reefs. In short, we followed an iterative process in which connected components were assigned the same label, thus being identified as forming the same component. Polygons were considered connected if they intersected (i.e., sharing a common boundary). To efficiently compute the intersections among polygons, we used the Sort-Tile-Recursive algorithm [29] implemented in Python Shapely library [30]. The implementation of the algorithm can be found in the Preprocessing.py module at [31]. The processing and all subsequent analyses were performed in a High Performance Cluster (HPC) consisting of 960 cores and 12TB of RAM.

Coral reefs less than 1000 m^2^ were considered potential noise in the dataset, leading us to disregard them. We made this choice based on the fact that the ACA is obtained from satellite imagery with about 3 m resolution. Thus, coral reefs less than 1000 m^2^ would correspond to fewer than 100 pixels. Furthermore, an inspection of the classified cover together with the satellite images suggested that many potential coral reefs (as per our definition) less than 1000 m^2^ were not accurately classified by the ACA (see Supplementary Fig. S1). Thus, including the classified polygons would result in a noncanonical dataset, which would bias the results (i.e., power-law exponents and fractality) obtained at this and smaller spatial scales.

### 2.3 Coral reef area, perimeter, and inter-reef distance

We computed coral reef area and perimeter using the *geopandas*.*GeoSeries*.*area* and *geopandas*.*GeoSeries*.*length* methods in geopandas Python’s library [32]. For each coral reef, we defined the inter-reef distance as the distance to its nearest neighbour. The inter-reef distance distribution was derived by calculating the distance to the nearest neighbor for each reef. To make this computation efficient, we used the Sort-Tile-Recursive algorithm [29] implemented in the Python Shapely library [30]. The implementation of the algorithm can be found in [31].

We note that our estimates of the total area for each coral reef province (Supplementary Table S2) slightly differ from that directly provided by the ACA because we removed reefs smaller than 10^3^ m^2^. Of course, if the area is computed before this data cleaning step, the results are identical (see Supplementary Table S1). The larger estimates of the coral habitat area provided by Lyons et al. [24] are based on the sum of the coral/algae and rock classes of the ACA, while our estimate is based only on the coral/algae class. Thus, our estimate can be understood as a lower bound of the coral habitat area.

### 2.4 Coral reef size distribution

We fitted the coral reef size data using the power-law package in Python [33, 34]. We performed goodness-of-fit tests using a range of alternative distribution models, including log-normal, exponential, and stretched exponential distributions. We found that the power-law distribution (including its truncated form, which in fact dominates) provided a significantly better fit to the data than any of the alternative models with *x*_min_ ranging from 10^3^ m^2^ to 10^4^ m^2^ (Supplementary Tables S3 and S4).

### 2.5 Box-Counting fractal dimension

Fractal dimensions are a generalisation of the concept of dimension to noninteger values and are used to measure the complexity of an object. Regular Euclidean objects have integer dimensions, with *D* = 1 for a line, *D* = 2 for a plane, and *D* = 3 for a volume. However, many natural objects are irregular and exhibit fractal dimensions between these integer values. For example, the dimension of the coastline of Great Britain is not 1, as one would expect from a smooth line, but around 1.25 [35]. The fractal dimension of an object can be computed using different methods, such as the Box-Counting algorithm.

We computed the Box-Counting fractal dimension of all mapped areas following a box-counting algorithm [35]. Briefly, the method computes the number of boxes of length *ϵ, N* (*ϵ*), needed to cover the underlying object. Then, the fractal dimension is simply defined as,

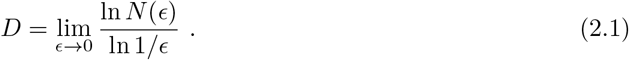

In practice, the mathematical limit *ϵ* → 0 is unreachable, and the fractal dimension is computed from the slope obtained in the plot of ln *N* (*ϵ*) versus ln(1*/ϵ*). To efficiently compute the number of overlapping boxes, we used the Sort-Tile-Recursive algorithm [29] implemented in the Python Shapely library [30]. The implementation of the algorithm can be found in [31].

### 2.6 Fractal dimensions from area-perimeter relation

The area of regular objects such as squares or circles scales as the square of the perimeter *A* ∼ *P* ^2^, while the area of irregular fractal objects scales more generally as *A* ∼ *P*^*σ*^, where *σ* = *D*_*A*_/*D*_*P*_ with *D*_*A*_ and *D*_*P*_ being the fractal dimension of the area and the perimeter, respectively [35, 36]. These fractal dimensions can be easily computed from the area-perimeter scaling exponent, *σ*, as *D*_*P*_ = (2 + *σ*)/2*σ* and *D*_*A*_ = (2 + *σ*)/2 [36].

The fractal dimension of the perimeter, *D*_*P*_, measures the complexity of the contour of the object, while the fractal dimension of the area, *D*_*A*_, measures the complexity of the object’s surface. The fractal dimension of the perimeter is related to the object’s roughness, with values closer to 1 indicating a smoother contour, and values closer to 2 indicating a more convoluted contour. On the other hand, the fractal dimension of the area is related to the space-filling properties of the object, with values closer to 2 indicating a more compact object and values closer to 1 indicating a poor space-filling object [35].

### 2.7 Compactness and elongation index

The compactness measurement is defined as the isoperimetric quotient,

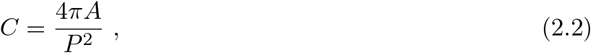

where *A* and *P* are the area and perimeter of the object under study, respectively. The compactness is a measure of how compact the object is, with values close to 1 indicating a more compact object and values close to 0 indicating a less compact object. Note that an object can have low compactness while being circular if it has holes in its surface (e.g., a doughnut).

The elongation index is defined as the Flaherty & Crumplin [37] length-width measure, stated as measure LW 7 in [38] and implemented in PySAL Python’s library [39]. The index is given by the minimum shape diameter (the shortest distance between two points on the object) divided by the maximum shape diameter (the longest distance between two points on the object). Values close to 1 indicate a more rounded object and values close to 0 a more elongated one.

## 3 Results

### 3.1 Coral reefs macroecological patterns

The area of all coral reefs within each province reported by the ACA was computed after processing the data to identify individual reefs (see Methods). We identified a total of 1, 579, 772 individual shallow-water tropical reefs (>1000 m^2^), which extend over a total of 52, 423 *km*^2^ of the ocean area. The comprehensive nature of this openly available dataset (see Methods), which includes all shallow-water tropical coral reefs worldwide allows the mean and median size of individual reefs to be estimated at 3.32 ha and 0.3 ha, respectively, across all coral reef provinces of the Atlas (see Supplementary Table S2 for statistics in each province).

Our analysis reveals that the size-frequency distribution of coral reefs converges to a power law distribution in all provinces, *y* ∼ *x*^−*α*^, which holds over multiple scales (i.e. from 100 km^2^ to 1000 m^2^), where *y* reflects the probability of occurrence of reefs of the area class *x km*^2^ (Fig. 1 a-b). This behaviour was consistent across all provinces within a range of 3 to 5 orders of magnitude in the coral reef area, yielding an average exponent of ⟨*α*⟩ = 1.84 (95% CI: 1.55 to 2.12) (see Methods). The global size-frequency distribution of coral reefs, fitted to all data, conforms to a power-law with an exponent of *α* = 1.8, which is consistent with the mean exponent ⟨*α*⟩. This provides evidence for a universal scaling law that governs the size distribution of coral reefs.

**Figure 1:**
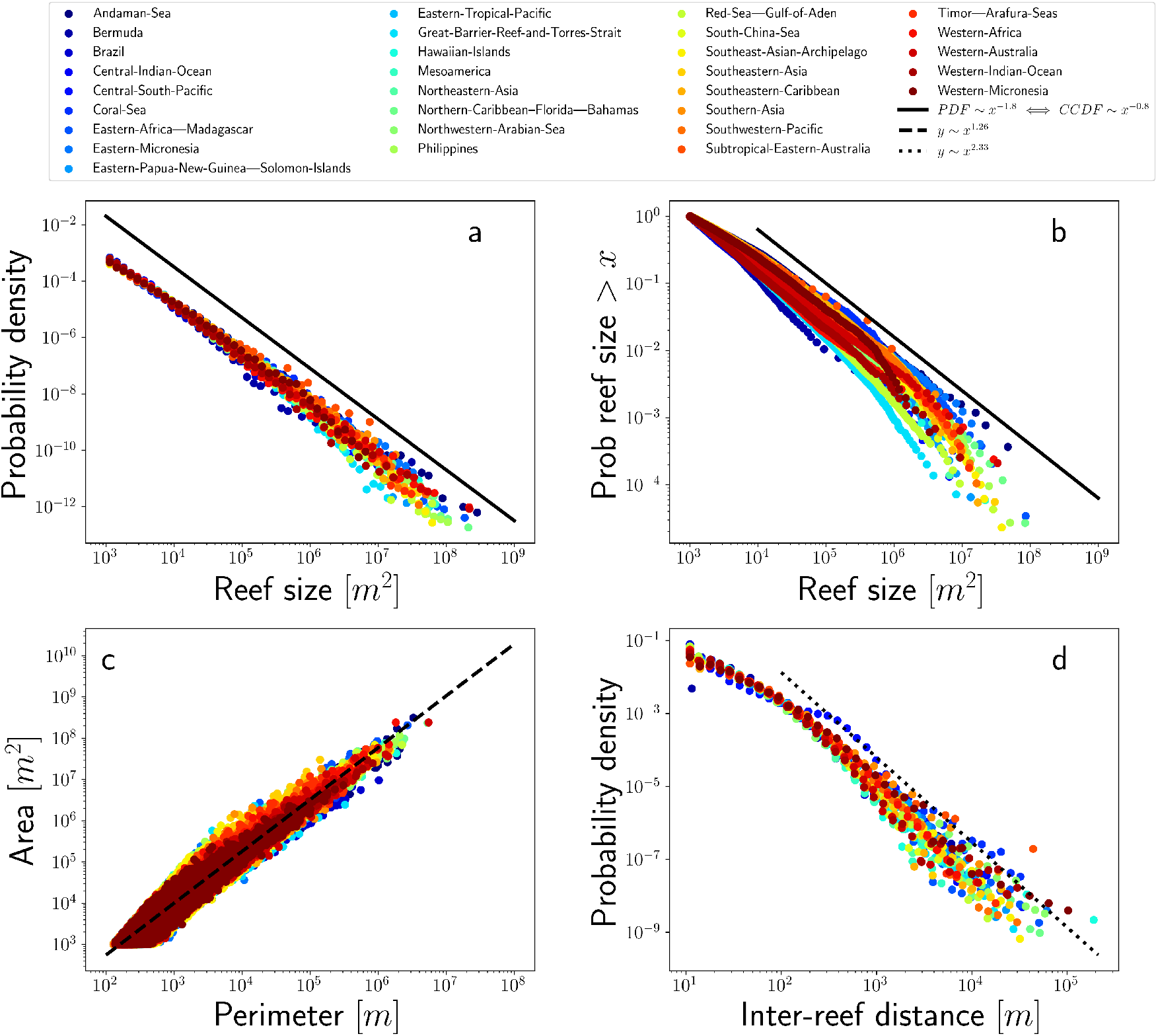
Macroecological patterns of global coral reef size, geometry, and spacing. (a) Size distribution. The black line corresponds to a fitted power-law of exponent 1.8. (b) Complementary cumulative distribution function (CCDF) with a corresponding exponent of 1.8. (c) Area-Perimeter relation. The black dashed line corresponds to a fitted power-law with an exponent of 1.23. (d) Inter-reef distance distribution. The black dotted line corresponds to a fitted power-law to the distribution tail with an exponent of 2.33.

The presence of a power-law size distribution also suggests that the object studied may be fractal in nature [40–44], at least along a certain range. A simple way to test whether coral reefs are fractals is to use the area-perimeter relation [35, 45], a method of fractal analysis that characterises the complexity of irregular shapes by examining the relationship between their area and perimeter (see Methods). The scaling of the coral reef area to the perimeter of all individual coral reefs also converges into a single power-law with an exponent of ⟨*α*⟩= 1.2578 (95% CI: 1.2573 to 1.2583), again indicating a universal behaviour (Fig. 1 c). When the relationship is fitted for each province independently, we found a mean exponent of ⟨*α*⟩ = 1.2574 (95% CI: 1.1757 to 1.3391), practically identical to the general exponent. From the area-perimeter scaling exponent, we can obtain the average fractal dimensions of the perimeter and area of the coral reefs under study (see Methods). We obtain an average perimeter fractal dimension of *D*_*P*_ = 1.2950 and a surface fractal dimension of *D*_*A*_ = 1.6289. Considering each province independently, the fractal dimensions range from 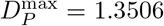 to 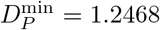 and from 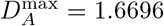 to 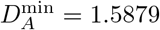 within a 95% confidence interval. This further confirms that the fractal dimensions are significantly different from the expected Euclidean dimensions of a line, *D*_*P*_ = 1, and a plane, *D*_*A*_ = 2. Thus, coral reefs develop fractal-like geometries, exhibiting complex, self-similar structures across different scales, which means that each portion can be considered a reduced-scale image of the whole [46].

The spatial distribution of coral reefs within each province was also investigated employing the inter-reef distance, defined as the minimum distance between a reef and its nearest neighbour. We found a heavy-tailed relation where the tail conforms to a power-law with an exponent of 2.33 (Fig. 1 d). This reveals that most of the reefs are close to each other at a distance of 10 m to 100 m, while few of them are separated from their nearest neighbour by more than 1 km, i.e. isolated. This finding is again mainly independent of the geographical location of the coral reefs studied, which arises as a universal property of the coral reef provinces. Furthermore, we computed the spatial autocorrelation of reef sizes, showing that this quantity is not randomly distributed (Supplementary Table S5).

### 3.2 The fractal nature of coral reefs

We computed the fractal dimension of the area and perimeter of each individual reef of all provinces using the box-counting algorithm (see Methods). The mean values for the fractal dimension of the perimeter, *D*_*P*_ = 1.24 (95% CI: 1.13 to 1.35) and the fractal dimension of the area, *D*_*A*_ = 1.60 (95% CI: 1.39 to 1.81), are well defined and consistent with those obtained from the area-perimeter relationship (Fig. 2 a,b). The fractal dimension of reef areas remains fairly consistent around the mean, though it shows a slight increase for larger coral reefs. (Fig. 2 a). However, the fractal dimension of the perimeter shows a more pronounced increase as a function of the size of the reef. This indicates that as coral reefs grow, their contour tends to become more and more convoluted, increasing their complexity, while their surface remains geometrically more stable.

**Figure 2:**
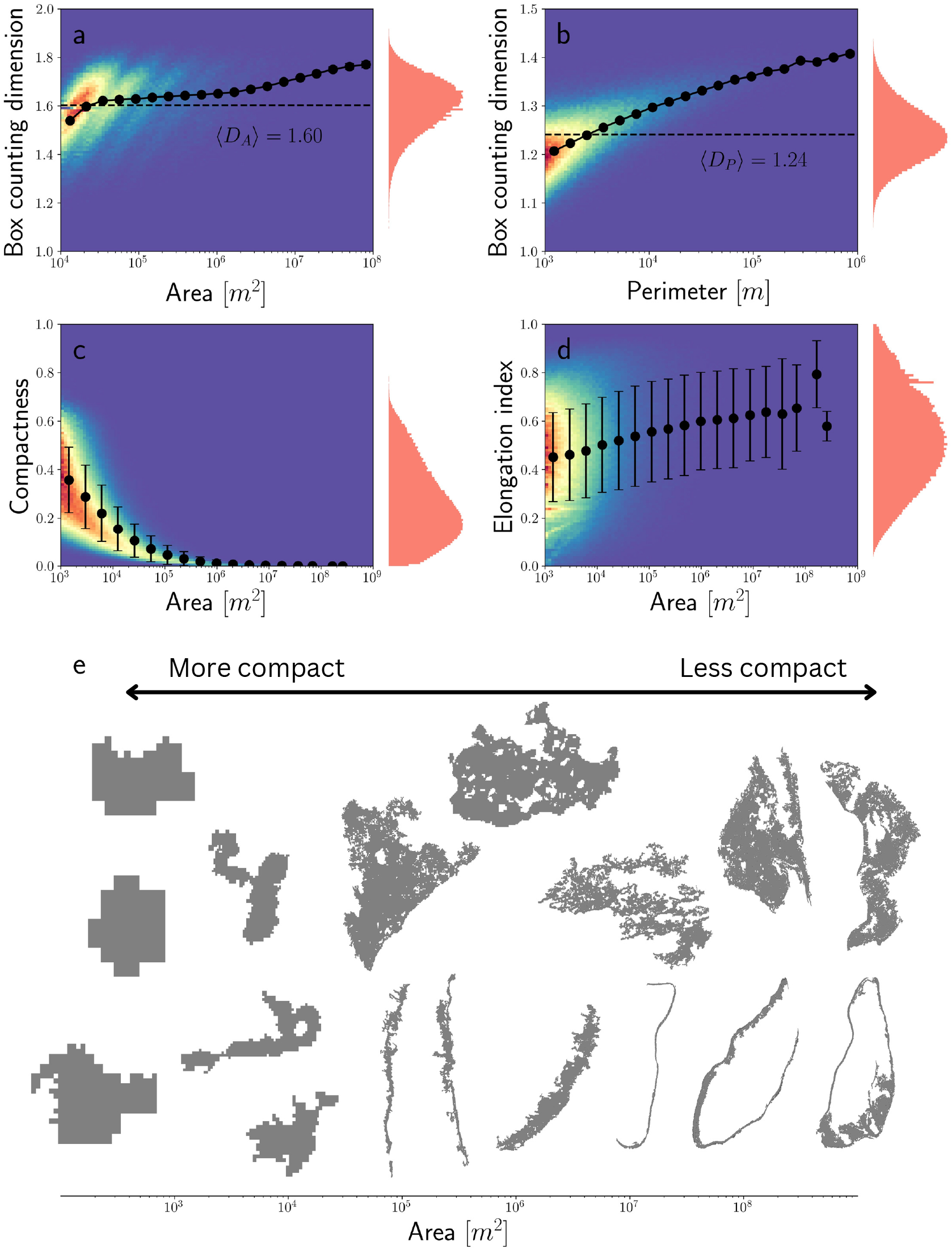
The fractal nature of global coral reefs. 2D histograms from all shallow-water coral reefs worldwide for: (a) the surface fractal dimension and area; (b) the perimeter fractal dimension and perimeter; (c) the compactness and area and (d) the elongation index and area. The black line corresponds to the mean values of the Y axis measure as a function of the X axis measure. The red histogram represents the distribution of the Y axis measure. (e) Example of coral reefs shape as a function of their surface.

To better understand the reef formation process from a geometric perspective, we computed other shape measurements such as compactness and elongation indices. We observe that the compactness of coral reefs decreases rapidly with increasing size (Fig. 2 c). Two changes of shape are consistent with this result: transitioning from round to elongated shapes or keeping their rounded shape but becoming more complex with discontinuities (holes), for example, sandy areas or reef lagoons, within the area mapped as coral/algae. These two processes are not necessarily mutually exclusive, as elongated shapes could appear from the evolution of empty, rounded shapes. For example, circular reefs could break their contour due to erosion or other hydrodynamic processes, giving rise to elongated shapes. The results obtained for the measurement of the elongation index suggest that both processes could occur, as the elongation of the reef increases with size while showing a very high variance. This hypothesis can be contrasted by examining the different shapes that coral reefs have formed.

In (Fig. 2 e) different coral reefs are located on the x-axis according to their size. We observe that they become less compact as they grow (as shown in panel Fig. 2 c), but this can occur by either keeping rounded shapes while becoming empty (first row of reefs in panel e) or by getting elongated (second row of reefs in panel e). Small coral reefs are mostly rounded and filled, whereas larger coral reefs are either elongated or have holes in their surface. Altogether, our analysis suggests that coral reefs tend to evolve from simple rounded filled shapes (high compactness and low elongation index) to more complex elongated and less compact forms (low compactness and high elongation index), giving rise to fractal objects with a stable surface fractal dimension and increasing perimeter fractal dimension as they grow (Fig. 2 e).

### 3.3 Fractality extends up to coral provinces

The fractal dimension of coral reefs varies across a range of spatial scales, spanning from individual colonies to entire reef systems [47]. This variability arises from the different physical, biological, and geological processes involved in the formation of structures present at each organisation level.

The provinces mapped by the ACA illustrate established patterns of coral reef biogeography with similar reef type and environmental conditions [15, 25], which we hypothesise can be understood as an upper organisational level for coral reefs. The processes involved in maintaining such large structures might be different from those of individual reefs, giving rise to a different fractal dimension. To investigate this hypothesis, we computed the surface fractal dimension for each coral province as a whole using the box-counting algorithm (Supplementary Fig. S2, Methods).

We determined a mean surface fractal dimension of ⟨*D*⟩ = 1.42 (95% CI: 1.24 to 1.59), with similar results across the different coral provinces under the assumption of a normal distribution (Fig. 3). Although this measure is consistent across all provinces, it is different from the mean surface fractal dimension of the individual reefs that make up the provinces. This observation suggests, once again, that coral reefs are self-organised systems that exhibit similar patterns of complexity and irregularity at different scales, and that coral reef provinces might be understood as the largest organisational level of coral reefs. Furthermore, we note that the fractal dimension of coral reef landscapes is similar to those anticipated from sizes expanding following a Fibonacci series, which yields a fractal dimension of 1.44 [48].

**Figure 3:**
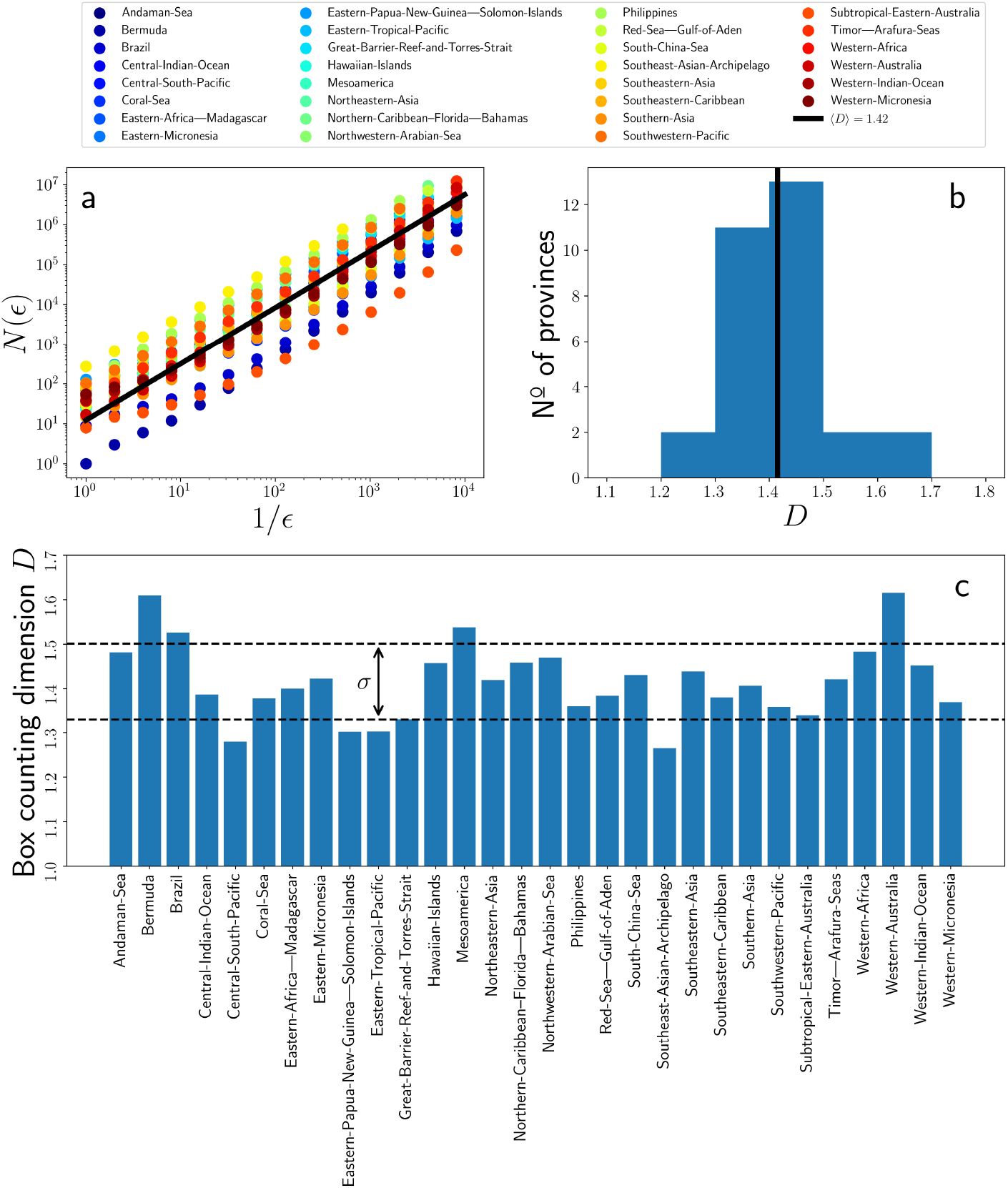
Box Counting Dimension of coral reef provinces surface. (a) Scaling of the measure (number of boxes of length *ϵ, N* (*ϵ*)) as a function of the ruler used (box length *ϵ*). The units of *ϵ* are degrees, with 1º being approximately 100 km. The slope of the fit for each province corresponds to its Box-Counting surface fractal dimension, *D*. (b) Histogram of the obtained Box-Counting fractal dimensions. The solid black line represents the mean Box-Counting fractal dimension, ⟨*D*⟩. (c) Box-counting surface dimension for each coral province. Black dashed lines correspond to a 1 *σ* deviation from the mean.

## 4 Discussion

Coral reefs self-organise to form macroecological patterns largely independent of their geographical location. This suggests that the local, short-term physical conditions experienced by each coral province have a limited effect on establishing these scaling laws. Coral reef structures exhibit complex fractal geometries with average fractal perimeter and surface dimensions of *D*_*P*_ = 1.24 and *D*_*A*_ = 1.60. The size-frequency distribution of these structures follows a power-law size distribution of exponent 1.8 and the inter-reef distance distribution is characterised by a heavy-tail that follows a power-law of exponent 2.33. Many reefs are relatively small, with an average surface of 3.32 ha, while few of them are massive, with surfaces reaching 100 km^2^. Similarly, most reefs are only 10 m to 100 m away from their nearest neighbour, with only a few being isolated, separated by more than 1 km. The universality of the observed patterns suggests that these features might likely result from the highly conserved interactions of biological, physical, and chemical processes, which, over geological timescales, lead to common reef landscape patterns across all provinces. Despite this, the precise mechanisms remain mostly unexplored, as we still lack mechanistic models that can generate coral reef landscapes across different scales and over time.

These universal scaling properties should be integrated into mechanistic models designed to describe coral reef development, which have previously lacked such constraints, whether considering individual reefs or entire coral reef provinces. For example, at the scale of a coral reef province, the inter-reef distance likely affects the interaction between coral reefs through hydrodynamic flows, influencing the dynamics of limiting resources transported to support photosynthesis and calcification. This process is further constrained by changes in sea level and the available vertical space [49–51]. This could also be linked to the formation of spur and groove structures, resulting from the circulation patterns of counter rotating circulation cells created by incoming surface waves, playing a pivotal role in shaping the complex topography of coral reefs [52]. Although our analysis is two-dimensional and focuses on the upper 10 m layer of the reefs supporting living coral/algae communities, spur and grove formations influence the dissipation of wave action [53], and may affect reef carbonate accretion-dissolution balances. Accurate models of coral reef growth and dynamics should replicate the universal features described here, including the power law distribution of coral reef sizes, fractal geometries, and size-dependent changes in reef shape.

Carbonate deposition is strongly influenced by biotic factors and, over geologic timescales, is further modulated by chemical processes that play a significant role in shaping reef architecture. [6]. Fractality in coral reef landscapes has been suggested to arise from multi-scale random processes that generate chaotic variability, influencing the population dynamics of frame-building corals and contributing to reef accretion [6]. Research indicates that modelled oscillations in population dynamics and growth result in fractality and power-law distributions in these landscapes [54–56]. While these models highlight the importance of growth processes, disturbance regimes promoting fragmentation are increasingly recognised as major drivers of complex landscape formation [57, 58]. Indeed, previous reports of regional fractality in Arabian Gulf coral reef landscapes have been explained as the outcome of cyclic disturbance regimes promoting fragmentation [59]. Dissolution processes, facilitated by the loss of the protective living biological cover that renders the carbonate framework vulnerable to dissolution, can also create complexity of carbonate reef structures at the landscape level. In addition, chemical (e.g., high CO_2_ and low pH) and physical (e.g., low temperature) conditions leading to thermodynamic forcing toward dissolution can favour these dissolution processes. Coral reefs typically accrete and extend in the fore-reef areas by receiving the flow of associated carbonate ions, minerals, and nutrients, and degrade in the back reef, where dissolution solutions prevail, leaving behind coral rubble [60]. Hence, the balance between biological-driven reef accretion and physical-chemical-driven erosion is not randomly distributed along the reefs and influences the dynamics of reefs over the millennial timescales of reef development, and hence the reef shape.

We have shown that the shape of coral reefs tends to vary with size and thereby during growth: from small, compact, and rather circular structures to big, complex forms with highly convoluted perimeters and low compactness. The monotonic decrease in compactness with increasing size can be attributed to either circular reefs developing multiple holes in their surface, a large inner lagoon, or becoming increasingly elongated. The development of elongated fringing reefs has been postulated to result mostly from reduced accommodation space as coral reefs grow and the interaction with sea level [60], but has also been explained as the result of the interaction between reefs and resources advected along hydrodynamic flows in a prevalent direction [50]. Reef patches inside reef lagoons can also be observed. As coral reefs expand in size, inner lagoons are formed and filled through various processes over multiple timescales [49, 61, 62]. Eventually, smaller reefs appear within these lagoons, starting as compact coral heads that eventually develop empty inner spaces on geological timescales. This phenomenology might suggest that a Turing instability [63, 64], a classical mechanism that leads to the formation of periodic patterns, is operating. In fact, this mechanism has been previously suggested [50], as a result of the interaction between nutrient diffusion and the processes of nutrient uptake and recycling within the reefs. However, the Turing mechanism would yield a normal distribution of inter-reef distances. This is not compatible with the heavy-tailed inter-reef distance distribution found in this study or with the observed power-law scaling. Thus, the Turing mechanism shows inconsistencies with the observed reef landscape patterns, although it may be in operation in certain instances.

The results discussed here are based on the analysis of remote sensing data from the ACA, which provides a comprehensive inventory of coral reefs worldwide. Nonetheless, there are some limitations to the ACA dataset that must be acknowledged [24, 65]. The ACA data are based on satellite imagery, which has a limited spatial resolution of 3 m, making it inadequate for resolving coral reef structures at finer scales. Moreover, reefs smaller than 1000 m^2^ were not consistently classified into habitat types, placing a minimum limit to the size of coral reefs included in our analyses. To conduct a proper analysis at these and lower spatial scales, much higher resolution would be necessary. This can be achieved using other instruments, such as drones, though achieving global coverage with these means is improbable. In addition, the ACA data encompasses only shallow-water tropical reefs and does not include deep-water reefs.

Another limitation of our results is that they are based on the coral/algae class of the ACA [15], which includes reefs where calcification processes are not necessarily dominated by corals but where other organisms, such as crustose, calcifying algae, and other encrusting calcifiers may have important contributions [66]. As a result, the growth/loss processes that lead to the universal power laws shown here do not necessarily result from the dynamics of scleractinian corals alone but, more broadly, from the calcifying community that lives in reefs as a whole. Unfortunately, the current structure of the living biological community in the shallow coral reefs assessed does not convey the historical balance between coral and non-coral calcifying organisms along the millennial timescales of reef formation. Thus, the analysed data reflect the current coral cover, and not the coral cover of the reefs during their extended formation process, much of which predate anthropogenic disturbance. Given the substantial coral cover loss during the last decades [27], it is likely that the present coral cover retrieved by the ACA underestimates the historic contribution of scleractinian corals to reef formation. Expanding our analysis to incorporate areas of reefs currently devoid of living coral/algae cover, by including the ACA’s rock class into the analysis, may shed light on the contribution of geological and biological processes to tropical reef landscapes, but also introduce noise derived from some of these rocky reefs not being biogenic in origin.

Another inherent limitation of our study is that some reefs may represent geological structures or structures produced by calcifying organisms other than corals. Some of the coral reefs in the dataset may have experienced significant erosion after coral loss, with hydrodynamic regimes possibly exerting a different forcing on reef degradation than on reef growth. Dynamic, mechanistic models are required to better understand the interplay between hydrodynamic regimes, coral cover, reef expansion, and erosion, ultimately leading to the universal patterns reported here. Finally, the ACA dataset is based on artificial intelligence algorithms that classify the benthic habitat using remote sensing data and therefore may be subject to errors in classification and interpretation [65]. Despite these limitations, the ACA dataset provides an unparalleled opportunity to study coral reef size and geometry on a global scale, with potentially useful applications.

Restoration strategies define quantitative targets, often expressed as a percentage of the area covered by coral reefs (e.g., 30%). To support these efforts, it is essential to quantify this area, a process that can be refined at the level of individual reefs using more detailed data. Our estimate of 52,423 km^2^ provides a lower bound for the area occupied by tropical shallow-water coral reefs, whereas the estimate of Lyons et al. of ∼ 80, 000 km^2^ can be viewed as an upper bound, as it includes both the coral/algae and rock classes of the ACA [24]. In any case, restoration efforts that reach or exceed hectare size must increase in the future to achieve the targets set by the Kunming-Montreal Global Biodiversity framework. Overall, the macroecological characterisation of universal laws governing the geometry of coral reefs, along with a comprehensive global coral reef dataset, represent a valuable resource for designing effective coral reef conservation and restoration projects, as well as optimise and quantify the effort and resources required.

We have identified different universal patterns that govern reef size and structure consistent across all tropical coral reef provinces worldwide. The observed patterns include a power-law size-frequency distribution with an exponent of 1.8, average perimeter and surface fractal dimensions of *D*_*P*_ = 1.24 and *D*_*A*_ = 1.60, respectively and an inter-reef distance distribution with a heavy-tail power-law with an exponent of 2.33. Despite the diverse biogeographical histories of the studied coral reefs, they exhibit similar spatial features across various scales and biotic and abiotic gradients. The mechanisms behind these phenomena remain unexplained, and uncovering them could provide crucial insights into the structure and function of coral reefs.

## Data availability statement

Data is publicly accessible in a Zenodo repository (https://doi.org/10.5281/zenodo.10025082) [22].

## Code availability statement

Code is publicly accessible in a Zenodo repository (https://doi.org/10.5281/zenodo.13952426) [31].

## Acknowledgements

A.G.R. and M.A.M. were supported through grant PID2021-123723OB-C22 (CYCLE) funded by MCIN/AEI/10.13039/501100011033 and by “ERDF A way of making Europe” and through grant CEX2021-001164-M (María de Maeztu Program for Units of Excellence in R&D) funded by MCIN/AEI/10.13039/501100011033. C.M.D. was funded by baseline funding provided by King Abdullah University of Science and Technology. We thank Sebastian Schmidt-Roach, Shannon Klein, Alexandra Steckbauer, Eleonora Re and Nayra Pluma for useful discussions in several meetings. Likewise, we acknowledge Miguel Álvarez-Alegría, Eva Llabrés, Damià Gomila, Tomàs Sintes, Emilio HernÁndez García, Pablo Moreno-Spiegelberg, and Núria MarbÀ for useful discussion. We also thank the team delivering the Allen Coral Atlas for the resource they made available, which enabled this research.

