## Supplementary Information for "Unravelling the universal spatial properties of coral reefs"

### 1 Allen Coral Atlas data

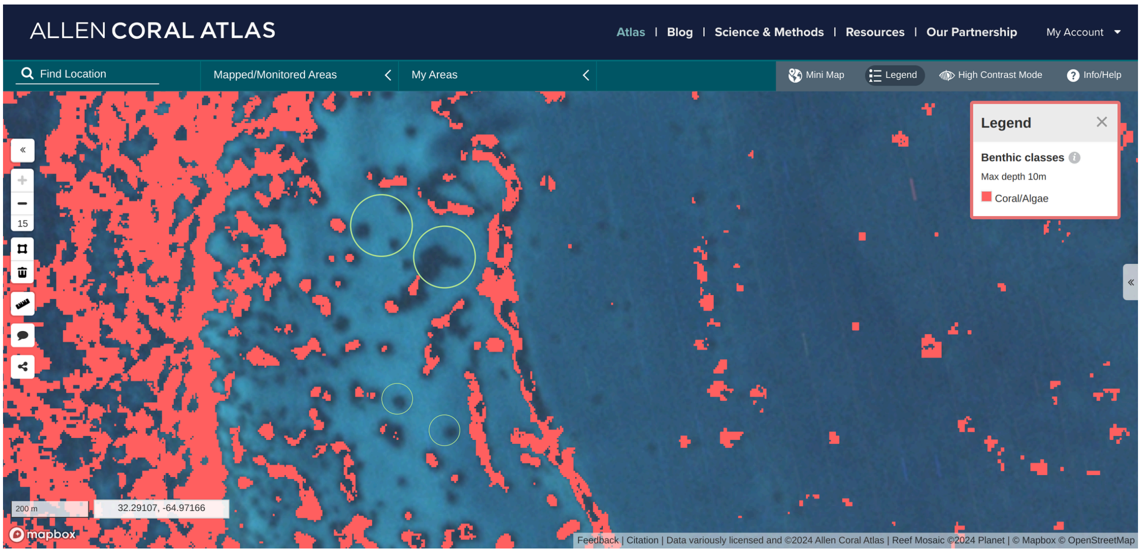


**Figure S1:** Example of coral/algae class polygons classified by the Allen Coral Atlas. The small polygons are not reliable for statistical analyses, as many might be missing (green circles).

**Table S1:** Comparison of the estimated total area for each coral reef province provided by the Allen Coral Atlas, and our work previous and after the data processing step.

| Province | Total Coral Area - Allen Coral Atlas (km²) | Total Coral Area - Before Processing (km²) | Total Coral Area – After Processing (km²) |
|---|---|---|---|
| Andaman-Sea | 2478.81 | 2478.81 | 2458.49 |
| Bermuda | 85.94 | 85.94 | 82.51 |
| Brazil | 129.12 | 129.12 | 117.61 |
| Central-Indian-Ocean | 957.7 | 957.7 | 937.56 |
| Central-South-Pacific | 529.41 | 529.41 | 509.03 |
| Coral-Sea | 210.61 | 210.61 | 204.5 |
| Eastern-Africa—Madagascar | 3206.32 | 3206.32 | 3168.68 |
| Eastern-Micronesia | 1460.07 | 1460.07 | 1445.35 |
| Eastern-Papua-New-Guinea—Solomon-Islands | 3721.66 | 3721.66 | 3643.85 |
| Eastern-Tropical-Pacific | 207.27 | 207.27 | 201.48 |
| Great-Barrier-Reef-and-Torres-Strait | 2103.55 | 2103.55 | 1952.24 |
| Hawaiian-Islands | 319.43 | 319.43 | 309.32 |
| Mesoamerica | 1672.81 | 1672.81 | 1644.13 |
| Northeastern-Asia | 445.72 | 445.72 | 438.56 |
| Northern-Caribbean–Florida—Bahamas | 4557.94 | 4557.94 | 4475.7 |
| Northwestern-Arabian-Sea | 910.71 | 910.71 | 873.91 |
| Philippines | 6319.9 | 6319.9 | 6231.33 |
| Red-Sea—Gulf-of-Aden | 3032.71 | 3032.71 | 2876.87 |
| South-China-Sea | 310.35 | 310.35 | 302.18 |
| Southeast-Asian-Archipelago | 9080.57 | 9080.57 | 8941.66 |
| Southeastern-Asia | 1508.35 | 1508.35 | 1490.35 |
| Southeastern-Caribbean | 1371.01 | 1371.01 | 1343.08 |
| Southern-Asia | 294.99 | 294.99 | 289.07 |
| Southwestern-Pacific | 3731.5 | 3731.5 | 3670.4 |
| Subtropical-Eastern-Australia | 32.29 | 32.29 | 31.9 |
| Timor—Arafura-Seas | 1783.06 | 1783.06 | 1743.77 |
| Western-Africa | 976.35 | 976.35 | 960.09 |
| Western-Australia | 1192.99 | 1192.99 | 1174.42 |
| Western-Indian-Ocean | 525.25 | 525.25 | 502.17 |
| Western-Micronesia | 415.86 | 415.86 | 403.21 |

### 2 Macroecological patterns

**Table S2:** Number of reefs, total reef area, total reef perimeter, mean reef area, mean reef perimeter and fractal dimension of coral reef provinces.

| Province | Nº reefs | Total reef area (km²) | Total reef perimeter (km) | Mean reef area (ha) | Mean reef perimeter (km) | Fractal dimension |
|---|---|---|---|---|---|---|
| Andaman-Sea | 2.46e+04 | 2458.49 | 82370.95 | 9.99 | 3.35 | 1.48 |
| Bermuda | 3.70e+03 | 82.51 | 4619.18 | 2.23 | 1.25 | 1.61 |
| Brazil | 5.20e+03 | 117.61 | 11040.94 | 2.26 | 2.12 | 1.53 |
| Central-Indian-Ocean | 3.11e+04 | 937.56 | 48889.55 | 3.02 | 1.57 | 1.39 |
| Central-South-Pacific | 3.04e+04 | 509.03 | 31947.16 | 1.68 | 1.05 | 1.28 |
| Coral-Sea | 5.53e+03 | 204.5 | 13578.79 | 3.7 | 2.46 | 1.38 |
| Eastern-Africa—Madagascar | 5.83e+04 | 3168.68 | 104343.91 | 5.44 | 1.79 | 1.4 |
| Eastern-Micronesia | 2.45e+04 | 1445.35 | 54452.86 | 5.91 | 2.23 | 1.42 |
| Eastern-Papua-New-Guinea—Solomon-Islands | 1.22e+05 | 3643.85 | 217711.52 | 2.98 | 1.78 | 1.3 |
| Eastern-Tropical-Pacific | 9.46e+03 | 201.48 | 10588.55 | 2.13 | 1.12 | 1.3 |
| Great-Barrier-Reef-and-Torres-Strait | 1.54e+05 | 1952.24 | 127829.58 | 1.27 | 0.83 | 1.33 |
| Hawaiian-Islands | 1.33e+04 | 309.32 | 17102.41 | 2.33 | 1.29 | 1.46 |
| Mesoamerica | 5.08e+04 | 1644.13 | 76462.11 | 3.24 | 1.51 | 1.54 |
| Northeastern-Asia | 1.44e+04 | 438.56 | 20612.65 | 3.05 | 1.43 | 1.42 |
| Northern-Caribbean–Florida—Bahamas | 1.12e+05 | 4475.7 | 179093.5 | 4.01 | 1.61 | 1.46 |
| Northwestern-Arabian-Sea | 3.13e+04 | 873.91 | 46168.12 | 2.79 | 1.47 | 1.47 |
| Philippines | 1.47e+05 | 6231.33 | 263475.43 | 4.25 | 1.8 | 1.36 |
| Red-Sea—Gulf-of-Aden | 1.64e+05 | 2876.87 | 186036.16 | 1.76 | 1.14 | 1.38 |
| South-China-Sea | 1.22e+04 | 302.18 | 16809.32 | 2.48 | 1.38 | 1.43 |
| Southeast-Asian-Archipelago | 2.59e+05 | 8941.66 | 426626.12 | 3.45 | 1.65 | 1.26 |
| Southeastern-Asia | 3.60e+04 | 1490.35 | 51462.3 | 4.14 | 1.43 | 1.44 |
| Southeastern-Caribbean | 3.67e+04 | 1343.08 | 71308.15 | 3.66 | 1.94 | 1.38 |
| Southern-Asia | 6.78e+03 | 289.07 | 13465.78 | 4.27 | 1.99 | 1.41 |
| Southwestern-Pacific | 9.54e+04 | 3670.4 | 164888.22 | 3.85 | 1.73 | 1.36 |
| Subtropical-Eastern-Australia | 5.66e+02 | 31.9 | 914.1 | 5.64 | 1.62 | 1.34 |
| Timor—Arafura-Seas | 5.02e+04 | 1743.77 | 77897.78 | 3.47 | 1.55 | 1.42 |
| Western-Africa | 2.09e+04 | 960.09 | 29485.25 | 4.6 | 1.41 | 1.48 |
| Western-Australia | 2.36e+04 | 1174.42 | 49788.67 | 4.97 | 2.11 | 1.62 |
| Western-Indian-Ocean | 2.36e+04 | 502.17 | 29976.67 | 2.13 | 1.27 | 1.45 |
| Western-Micronesia | 1.47e+04 | 403.21 | 24713.6 | 2.73 | 1.68 | 1.37 |

**Table S3:** Statistical comparison of power-law fit of the coral reef size distribution to other plausible distributions such as Exponential, Stretched exponential, Lognormal, Lognormal positive or Truncated power-law. A negative ratio implies that the compared distribution is more plausible than the assumed power-law, while a positive ratio indicates that the assumed power-law distribution is a better fit. The p-value measures the statistical significance of the result of each comparison, i.e. it must be $< 0.05$ for the result to be significant.

| | | Exponential | Stretched Exponential | Lognormal | Lognormal Positive | Truncated Power Law |
|---|---|---|---|---|---|---|
| **Andaman-Sea** | p-value | 1.24e-23 | 2.93e-03 | 6.21e-01 | 2.70e-02 | 6.03e-02 |
| | Ratio | 7958.48 | 24.72 | -0.40 | 13.52 | -1.76 |
| **Bermuda** | p-value | 4.79e-05 | 7.81e-03 | 8.64e-01 | 1.51e-02 | 9.96e-01 |
| | Ratio | 2768.75 | 14.95 | -0.01 | 12.44 | 0.00 |
| **Brazil** | p-value | 6.00e-31 | 1.73e-04 | 4.95e-01 | 3.31e-04 | 2.44e-01 |
| | Ratio | 3824.34 | 22.83 | -0.38 | 19.27 | -0.68 |
| **Central-Indian-Ocean** | p-value | 5.49e-151 | 1.65e-02 | 1.47e-08 | 6.27e-04 | 0.00e+00 |
| | Ratio | 17443.86 | -29.98 | -41.40 | -35.42 | -63.45 |
| **Central-South-Pacific** | p-value | 5.02e-62 | 8.63e-02 | 1.87e-01 | 2.72e-01 | 6.31e-07 |
| | Ratio | 3649.17 | 10.17 | -1.98 | 5.02 | -12.41 |
| **Coral-Sea** | p-value | 1.98e-98 | 5.67e-04 | 5.65e-01 | 5.47e-04 | 1.50e-07 |
| | Ratio | 4541.59 | 20.22 | -0.31 | 17.95 | -13.79 |
| **Eastern-Africa—Madagascar** | p-value | 9.13e-91 | 1.72e-25 | 6.28e-02 | 7.04e-20 | 7.31e-03 |
| | Ratio | 40940.17 | 201.41 | 0.33 | 143.46 | -3.60 |
| **Eastern-Micronesia** | p-value | 1.27e-80 | 4.21e-05 | 9.93e-01 | 1.46e-03 | 9.47e-03 |
| | Ratio | 5767.13 | 28.85 | 0.00 | 16.84 | -3.37 |
| **Eastern-Papua-New-Guinea—Solomon-Islands** | p-value | 6.01e-09 | 3.65e-01 | 1.62e-01 | 1.62e-01 | 6.97e-03 |
| | Ratio | 231.98 | -2.48 | -2.68 | -2.68 | -3.64 |
| **Eastern-Tropical-Pacific** | p-value | 1.57e-13 | 9.32e-01 | 1.42e-01 | 3.59e-01 | 1.19e-02 |
| | Ratio | 1655.07 | -0.42 | -3.42 | -2.98 | -3.16 |
| **Great-Barrier-Reef-and-Torres-Strait** | p-value | 1.11e-29 | 1.98e-01 | 1.14e-04 | 1.98e-01 | 9.99e-10 |
| | Ratio | 16943.45 | 20.29 | -23.03 | -13.15 | -18.66 |
| **Hawaiian-Islands** | p-value | 1.19e-12 | 1.29e-01 | 1.31e-01 | 4.65e-01 | 4.88e-03 |
| | Ratio | 6041.76 | 12.22 | -3.46 | 4.49 | -3.96 |
| **Mesoamerica** | p-value | 3.43e-75 | 2.80e-09 | 5.84e-01 | 1.04e-06 | 9.24e-06 |
| | Ratio | 16356.74 | 71.41 | -0.40 | 46.78 | -9.83 |
| **Northeastern-Asia** | p-value | 3.38e-71 | 4.70e-04 | 1.68e-06 | 3.30e-06 | 0.00e+00 |
| | Ratio | 6360.31 | -29.02 | -29.21 | -29.19 | -41.23 |
| **Northern-Caribbean–Florida—Bahamas** | p-value | 6.32e-69 | 3.44e-14 | 8.90e-01 | 7.65e-09 | 9.09e-03 |
| | Ratio | 29091.42 | 135.67 | -0.04 | 72.55 | -3.40 |
| **Northwestern-Arabian-Sea** | p-value | 3.91e-78 | 2.60e-17 | 8.19e-01 | 9.45e-16 | 4.98e-02 |
| | Ratio | 48828.62 | 183.48 | 0.06 | 162.94 | -1.92 |
| **Philippines** | p-value | 1.39e-286 | 1.26e-13 | 1.82e-05 | 8.97e-07 | 0.00e+00 |
| | Ratio | 71195.32 | 188.54 | -24.29 | 97.32 | -77.33 |
| **Red-Sea—Gulf-of-Aden** | p-value | 4.17e-42 | 2.35e-01 | 9.45e-04 | 2.20e-01 | 2.37e-06 |
| | Ratio | 15470.31 | 17.44 | -18.59 | -11.52 | -11.13 |
| **South-China-Sea** | p-value | 1.61e-76 | 3.76e-06 | 8.31e-01 | 4.34e-05 | 6.40e-04 |
| | Ratio | 7335.20 | 36.79 | -0.06 | 28.19 | -5.83 |
| **Southeast-Asian-Archipelago** | p-value | 0.00e+00 | 8.93e-02 | 3.49e-30 | 1.11e-12 | 0.00e+00 |
| | Ratio | 88730.10 | -54.99 | -182.78 | -158.78 | -215.45 |
| **Southeastern-Asia** | p-value | 1.59e-78 | 2.63e-01 | 1.50e-02 | 8.40e-01 | 4.98e-11 |
| | Ratio | 6729.44 | 9.15 | -6.99 | -1.16 | -21.59 |
| **Southeastern-Caribbean** | p-value | 3.46e-67 | 1.01e-04 | 2.21e-06 | 2.21e-06 | 0.00e+00 |
| | Ratio | 4468.22 | -29.16 | -28.16 | -28.16 | -40.67 |
| **Southern-Asia** | p-value | 8.59e-26 | 1.00e-01 | 5.96e-01 | 2.37e-01 | 5.18e-02 |
| | Ratio | 2295.95 | 7.27 | -0.39 | 4.18 | -1.89 |
| **Southwestern-Pacific** | p-value | 0.00e+00 | 2.38e-46 | 5.12e-01 | 1.72e-43 | 0.00e+00 |
| | Ratio | 103608.60 | 387.77 | -0.57 | 360.89 | -43.43 |
| **Subtropical-Eastern-Australia** | p-value | 8.79e-11 | 9.90e-01 | 3.70e-01 | 9.48e-01 | 4.65e-02 |
| | Ratio | 1047.13 | 0.03 | -1.08 | 0.17 | -1.98 |
| **Timor—Arafura-Seas** | p-value | 1.70e-97 | 1.89e-07 | 2.10e-01 | 2.46e-05 | 3.22e-07 |
| | Ratio | 20686.45 | 70.02 | -2.18 | 46.00 | -13.06 |
| **Western-Africa** | p-value | 1.45e-14 | 2.66e-09 | 2.64e-01 | 8.27e-07 | 9.70e-01 |
| | Ratio | 19217.88 | 97.00 | 0.10 | 58.74 | -0.00 |
| **Western-Australia** | p-value | 2.29e-15 | 6.49e-07 | 6.90e-01 | 1.80e-05 | 2.90e-01 |
| | Ratio | 14180.60 | 57.94 | 0.04 | 38.93 | -0.56 |
| **Western-Indian-Ocean** | p-value | 1.57e-43 | 3.21e-11 | 7.65e-01 | 7.13e-10 | 5.82e-02 |
| | Ratio | 22068.17 | 98.19 | 0.05 | 79.29 | -1.79 |
| **Western-Micronesia** | p-value | 1.12e-41 | 3.45e-03 | 1.00e-03 | 1.00e-03 | 4.14e-12 |
| | Ratio | 1432.37 | -13.38 | -12.08 | -12.08 | -24.03 |

**Table S4:** Results of the power-law fit of the coral reef size distribution. $\alpha$ is the exponent of the distribution, $D$ is the Kolmogorov-Smirnov distance (fit error), $\sigma$ is the standard error of the exponent, $x_{\max}$ is the optimal maximum value for the power-law fit and $x_{min}$ is the optimal minimum value for the power-law fit

| | $\alpha$ | $D$ | $\sigma$ | $x_{\max}$ | $x_{\min}$ |
|---|---|---|---|---|---|
| **Eastern-Micronesia** | 1.80 | 1.60e-02 | 1.23e-02 | 3.58e+07 | 1.76e+04 |
| **Southeast-Asian-Archipelago** | 1.78 | 1.22e-02 | 2.72e-03 | 6.93e+07 | 6.95e+03 |
| **Coral-Sea** | 1.65 | 2.48e-02 | 1.16e-02 | 6.34e+06 | 2.05e+03 |
| **Northwestern-Arabian-Sea** | 1.83 | 8.60e-03 | 5.11e-03 | 6.11e+07 | 1.25e+03 |
| **Western-Indian-Ocean** | 1.86 | 8.58e-03 | 7.21e-03 | 2.97e+07 | 1.88e+03 |
| **Southeastern-Asia** | 1.77 | 8.28e-03 | 9.57e-03 | 4.29e+07 | 1.47e+04 |
| **Northern-Caribbean–Florida—Bahamas** | 1.86 | 5.43e-03 | 6.15e-03 | 2.35e+08 | 1.21e+04 |
| **Hawaiian-Islands** | 1.84 | 8.31e-03 | 1.19e-02 | 4.34e+07 | 4.12e+03 |
| **Great-Barrier-Reef-and-Torres-Strait** | 1.98 | 8.10e-03 | 6.68e-03 | 6.27e+07 | 1.01e+04 |
| **Red-Sea—Gulf-of-Aden** | 1.95 | 1.02e-02 | 7.27e-03 | 6.92e+07 | 1.63e+04 |
| **Mesoamerica** | 1.83 | 7.59e-03 | 7.28e-03 | 7.67e+07 | 7.70e+03 |
| **Subtropical-Eastern-Australia** | 1.61 | 2.62e-02 | 2.55e-02 | 8.35e+06 | 1.01e+03 |
| **Southeastern-Caribbean** | 1.81 | 1.80e-02 | 1.03e-02 | 2.63e+07 | 2.00e+04 |
| **Central-South-Pacific** | 1.86 | 1.24e-02 | 1.29e-02 | 8.58e+06 | 1.02e+04 |
| **Western-Africa** | 1.93 | 1.18e-02 | 1.00e-02 | 2.40e+08 | 3.64e+03 |
| **Bermuda** | 2.06 | 1.95e-02 | 2.77e-02 | 3.44e+07 | 3.18e+03 |
| **Andaman-Sea** | 1.78 | 8.19e-03 | 1.14e-02 | 3.16e+08 | 1.89e+04 |
| **Western-Australia** | 1.83 | 1.00e-02 | 9.50e-03 | 2.42e+08 | 5.51e+03 |
| **Central-Indian-Ocean** | 1.71 | 1.27e-02 | 5.83e-03 | 1.37e+07 | 3.11e+03 |
| **Timor—Arafura-Seas** | 1.79 | 7.21e-03 | 6.47e-03 | 5.41e+07 | 5.88e+03 |
| **Brazil** | 1.82 | 1.39e-02 | 1.66e-02 | 9.54e+06 | 2.27e+03 |
| **Eastern-Tropical-Pacific** | 1.93 | 1.64e-02 | 2.04e-02 | 1.02e+07 | 1.06e+04 |
| **South-China-Sea** | 1.82 | 1.47e-02 | 1.06e-02 | 1.02e+07 | 3.21e+03 |
| **Northeastern-Asia** | 1.72 | 1.86e-02 | 8.95e-03 | 1.27e+07 | 4.10e+03 |
| **Eastern-Africa—Madagascar** | 1.79 | 1.23e-02 | 5.36e-03 | 2.07e+08 | 4.70e+03 |
| **Western-Micronesia** | 1.82 | 1.89e-02 | 1.65e-02 | 7.28e+06 | 1.70e+04 |
| **Southern-Asia** | 1.78 | 1.44e-02 | 1.90e-02 | 1.99e+07 | 8.45e+03 |
| **Philippines** | 1.75 | 5.20e-03 | 3.32e-03 | 1.17e+08 | 5.41e+03 |
| **Southwestern-Pacific** | 1.74 | 7.58e-03 | 3.01e-03 | 5.55e+07 | 2.05e+03 |
| **Eastern-Papua-New-Guinea—Solomon-Islands** | 2.41 | 2.20e-02 | 4.28e-02 | 2.92e+07 | 5.38e+05 |

### 3 Spatial autocorrelation of coral reef size

**Table S5:** Spatial autocorrelation analysis of the coral reef size distribution in different provinces. Moran's I is the measure of spatial autocorrelation of the distribution, Expected I is the expected value of Moran's I under spatial randomness and p-value is the significance of the result. A significant p-value (e.g. $p < 0.01$) indicates that the distribution is not spatially random.

| **Province** | **Moran I** | **Expected I** | **p-value** |
|---|---|---|---|
| Andaman-Sea | 3.92e-02 | -4.07e-05 | 1.00e-03 |
| Bermuda | 8.06e-02 | -2.70e-04 | 1.00e-03 |
| Brazil | 5.44e-02 | -1.92e-04 | 1.00e-03 |
| Central-Indian-Ocean | 3.12e-02 | -3.22e-05 | 1.00e-03 |
| Central-South-Pacific | 6.53e-02 | -3.29e-05 | 1.00e-03 |
| Coral-Sea | 2.40e-02 | -1.81e-04 | 1.00e-03 |
| Eastern-Africa—Madagascar | 1.51e-02 | -1.72e-05 | 1.00e-03 |
| Eastern-Micronesia | 7.17e-02 | -4.09e-05 | 1.00e-03 |
| Eastern-Papua-New-Guinea—Solomon-Islands | 5.35e-02 | -8.17e-06 | 1.00e-03 |
| Eastern-Tropical-Pacific | 6.15e-02 | -1.06e-04 | 1.00e-03 |
| Great-Barrier-Reef-and-Torres-Strait | 3.53e-02 | -6.51e-06 | 1.00e-03 |
| Hawaiian-Islands | 4.71e-02 | -7.55e-05 | 1.00e-03 |
| Mesoamerica | 6.15e-02 | -1.97e-05 | 1.00e-03 |
| Northeastern-Asia | 4.01e-02 | -6.95e-05 | 1.00e-03 |
| Northern-Caribbean–Florida—Bahamas | 2.76e-02 | -8.96e-06 | 1.00e-03 |
| Northwestern-Arabian-Sea | 3.20e-02 | -3.19e-05 | 1.00e-03 |
| Philippines | 6.22e-02 | -6.82e-06 | 1.00e-03 |
| Red-Sea—Gulf-of-Aden | 4.09e-02 | -6.11e-06 | 1.00e-03 |
| South-China-Sea | 3.09e-02 | -8.21e-05 | 1.00e-03 |
| Southeast-Asian-Archipelago | 5.84e-02 | -3.86e-06 | 1.00e-03 |
| Southeastern-Asia | 3.39e-02 | -2.78e-05 | 1.00e-03 |
| Southeastern-Caribbean | 2.24e-02 | -2.72e-05 | 1.00e-03 |
| Southern-Asia | 3.60e-02 | -1.48e-04 | 1.00e-03 |
| Southwestern-Pacific | 6.09e-02 | -1.05e-05 | 1.00e-03 |
| Subtropical-Eastern-Australia | 2.85e-02 | -1.77e-03 | 1.05e-01 |
| Timor—Arafura-Seas | 2.86e-02 | -1.99e-05 | 1.00e-03 |
| Western-Africa | 9.72e-02 | -4.79e-05 | 1.00e-03 |
| Western-Australia | 3.74e-02 | -4.23e-05 | 1.00e-03 |
| Western-Indian-Ocean | 6.36e-02 | -4.24e-05 | 1.00e-03 |
| Western-Micronesia | 3.16e-02 | -6.78e-05 | 1.00e-03 |

### 4 Box counting dimension of coral reef provinces

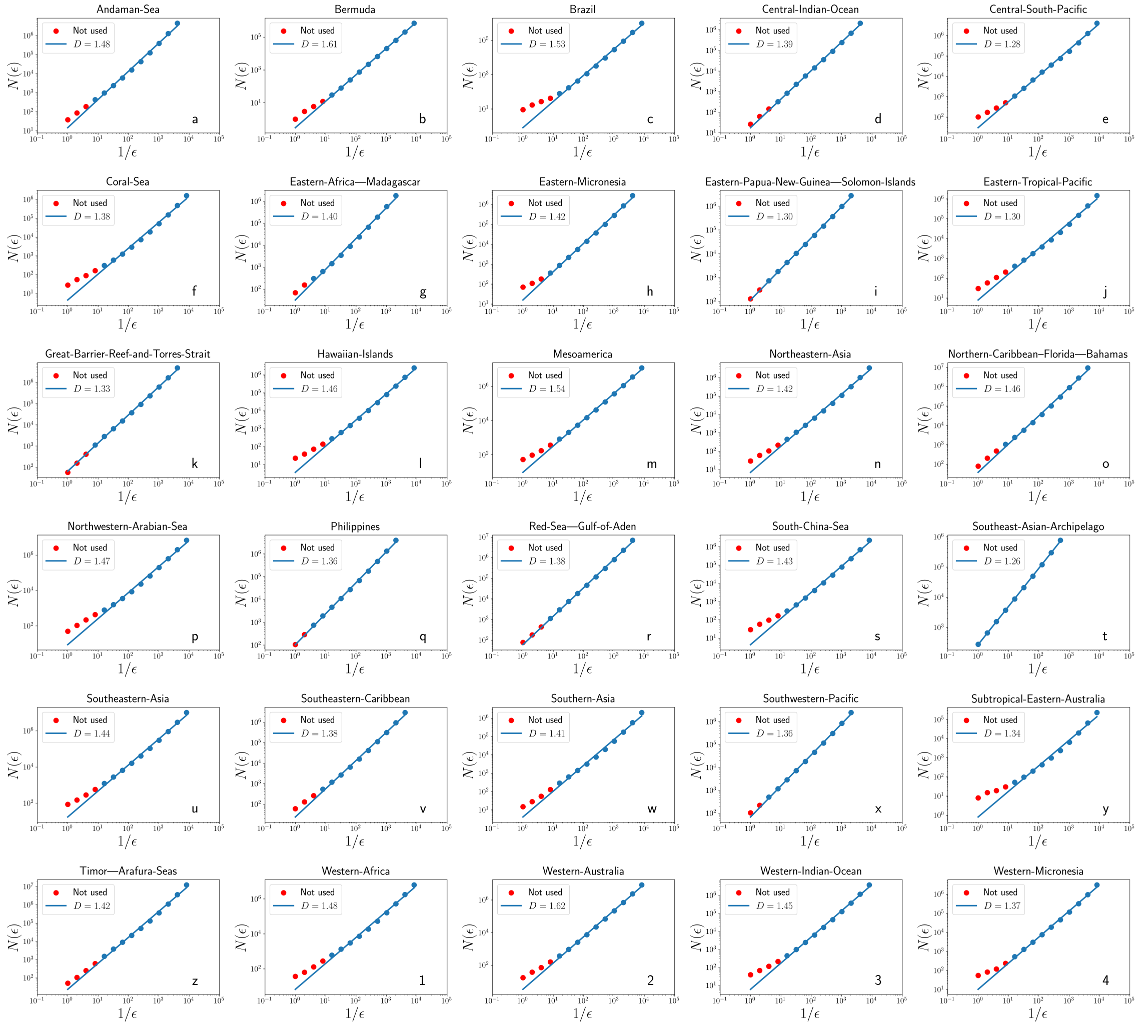


**Figure S2:** Computation of the fractal dimensions of each coral province using the box-counting method. $N(\epsilon)$ corresponds to the number of boxes of length $\epsilon$ needed to cover the object. The slope of $\log(N(\epsilon))$ vs $1/\epsilon$ approximates the fractal dimension of the object. Blue points were used to perform this computation while red dots were discarded.
